# Alcohol-Induced Mucociliary Dysfunction: Role of Defective CFTR Channel Function

**DOI:** 10.1101/2023.07.17.548927

**Authors:** Lawrence Rasmussen, Denise Stafford, Jennifer LaFontaine, Antonio Allen, Linto Antony, Hyunki Kim, S. Vamsee Raju

## Abstract

Excessive alcohol use is thought to increase the risk of respiratory infections by impairing mucociliary clearance (MCC). In this study, we investigate the hypothesis that alcohol reduces the function of CFTR, the protein that is defective in individuals with cystic fibrosis, thus altering mucus properties to impair MCC and the airway’s defense against inhaled pathogens.

**Methods:** Sprague Dawley rats with wild type CFTR (+/+), matched for age and sex, were administered either a Lieber-DeCarli alcohol diet or a control diet with the same number of calories for eight weeks. CFTR activity was measured using nasal potential difference (NPD) assay and Ussing chamber electrophysiology of tracheal tissue samples. In vivo MCC was determined by measuring the radiographic clearance of inhaled Tc99 particles and the depth of the airway periciliary liquid (PCL) and mucus transport rate in excised trachea using micro-optical coherence tomography (µOCT). The levels of rat lung MUC5b and CFTR were estimated by protein and mRNA analysis.

**Results:** Alcohol diet was found to decrease CFTR ion transport in the nasal and tracheal epithelium *in vivo* and *ex vivo*. This decrease in activity was also reflected in partially reduced full-length CFTR protein levels but not, in mRNA copies, in the lungs of rats. Furthermore, alcohol-fed rats showed a significant decrease in MCC after 8 weeks of alcohol consumption. The trachea from these rats also showed reduced PCL depth, indicating a decrease in mucosal surface hydration that was reflected in delayed mucus transport. Diminished MCC rate was also likely due to the elevated MUC5b expression in alcohol-fed rat lungs.

**Conclusions:** Excessive alcohol use can decrease the expression and activity of CFTR channels, leading to reduced airway surface hydration and impaired mucus clearance. This suggests that CFTR dysfunction plays a role in the compromised lung defense against respiratory pathogens in individuals who drink alcohol excessively.

**What is the current scientific knowledge on this subject?:** Excessive alcohol is associated with delayed mucociliary clearance (MCC) and an increased risk of respiratory infections among heavy drinkers.

**What does this study add to the field?:** Chronic alcohol use reduces CFTR activity and airway surface hydration explaining the mechanisms underlying mucociliary dysfunction. Acquired CFTR dysfunction may be a suitable target to improve host immunity in those affected by prolonged alcohol use.

## INTRODUCTION

Over 70% of adults in the U.S. who are 18 or older consume alcohol.^1–3^ Various pulmonary infections^17, 18^ are linked to the negative effects of excessive alcohol consumption, which is defined as more than 8 drinks per day.^2, 13–16^ Excessive use of alcohol increases the risk and severity of bacterial infections such as Klebsiella pneumonia, Streptococcus pneumonia, Legionella pneumophilia, Listeria monocytigenes, and Mycobacterium tuberculosis.^17, 18, 24^ Thus, more research is needed to gain insights into the mechanisms by which alcohol impairs lung host defense and develop potential interventions to reduce this risk.

Mucociliary clearance (MCC) is the process by which the mucus on the airway surface is moved upward and out of the lungs. This is a continuous process important for removing inhaled dust, bacteria, and other potentially harmful substances from the lungs. Alcohol use can have a negative effect on MCC, which can extend the retention time of pathogens and cause respiratory infections.^43^ Since therapeutic interventions that augment MCC are proven to reduce infections in individuals with chronic airway disease, restoring MCC to its physiologic levels in alcohol users may allow for effective removal of pathogens, preventing colonization and infection of lungs.^46^

MCC relies on the properties of two surface liquid layers: airway surface liquid (ASL) and Periciliary liquid (PCL). ASL is the outer liquid layer consisting of a highly viscous mucous blanket that traps microbes and dust. PCL is a thin, inner liquid layer that covers and lubricates cilia that beat in coordinated waves to create currents that propel mucus, which is expelled by swallowing/coughing. Research on the effects of alcohol on ciliary function has shown that prolonged alcohol exposure (>6 hours) significantly decreases ciliary beat frequency (CBF)^51–54^ and increases mucus expression compounding the burden on cilia.^55, 56^ However, there are currently no effective drugs to reverse ciliary defects to improve MCC in alcohol users. Hence, we explored the effects of alcohol on mechanisms involved in regulating airway surface hydration.

The PCL depth is physiologically maintained at ≥ 7 µm, the height of outstretched cilia, to support their beating. When the PCL height is at its maximum, mucus transport is at its highest.^49, 57^ PCL is critically regulated by the balance between the total number of Cl^-^ ions secreted by CFTR channels or Na^+^ ions absorbed by ENaC channels in airway epithelial cells.[1] In turn, Ion transport determines water transport across the epithelial cells available for PCL hydration. In vitro studies by us and others have shown that excessive alcohol consumption impairs the ability of CFTR channels to transport chloride and bicarbonate ions across the cell membrane.[2, 3] CFTR stands for cystic fibrosis transmembrane conductance regulator. In people with cystic fibrosis, the CFTR protein is defective, which leads to thick, sticky mucus that can clog the airways and cause respiratory infections.

So far, the effects of alcohol on CFTR are not validated in an animal model, nor whether loss of CFTR activity translates into decreased fluid transport across the airway surface to impact MCC are tested sufficiently. Therefore, we have used a rat model of alcohol administration to test the hypothesis that *alcohol reduces CFTR function and disrupts physiologic airway surface hydration and mucociliary transport*.

## METHODS

### Animal studies

All animal protocols were reviewed and approved by the University of Alabama at Birmingham Institutional Animal Care and Use Committee.

8-week-old Sprague-Dawley rats were obtained from Charles River Laboratories and allowed to acclimate for a week. Rats were housed individually in standard cages with bedding in temperature, humidity and light-controlled rooms with 12-hour light-dark cycles. Isocaloric (1 kcal/ml) control and ethanol liquid diets (Bio-Serv, Frenchtown, NJ) formulated according to Lieber and DeCarli were used for pair-feeding).[4, 5] The alcoholic diet contained 36% of total calories from ethanol, 11% carbohydrate, 18% protein, and 35% as fat, weight-matched controls received an isocaloric amount of diet in which ethanol calories were replaced by maltose-dextrins. Rats in the alcohol-fed group were acclimated to the diet by gradually increasing the concentration of alcohol in the diet from 0 to 3% (w/v) over 2 weeks and maintained at this level for 8 weeks until final assessments. The amount of liquid diet consumed by each ethanol-fed rat was measured for a 24-hour period, and each corresponding control rat was fed an equal amount of the control liquid diet during the next 24hr period.

### Radiographic in vivo mucociliary clearance assay

Rats were anesthetized in a controlled chamber containing isoflurane gas (5%, flow rate 3 L/min for 5 min). Adequate depth of anesthesia was ascertained both by observing decreased breathing rate and by no response to toe pinch. Once sedated, rats were placed behind a radioactive barrier and suspended by their incisors on an inverted plexiglass stand. Blunted forceps were used to secure the tongue without causing tissue damage. The tongue was pulled in the radial direction to obtain a clear view of the trachea. After clear visualization of the larynx, the trachea was intubated with a fiber optic bronchoscope adapted for use in rats. A flexible plastic tube modified from an Excel Safelet IV catheter was used to intubate the rats. The intubated rat was then attached to a Raindrop Nebulizer under isoflurane (2%, flow rate 0.5 L/min) to receive 99mTc-DPTA (10 mCi in 500µL PBS) aerosols. The nebulizer setup included a dosimetry system and a compressed air supply. After radioactive delivery, the rats were extubated and placed back into the anesthesia chamber at (3%, flow rate 2 L/min) and transferred for imaging. A gamma camera was used to measure the clearance of 99mTc-DPTA from rat lungs. Regions of the lungs were marked by a coincident CT image, and time-dependent clearance of radiolabel from the lungs over 30 minutes was calculated. The radioactive counts were corrected for decay and expressed as a percentage of radioactivity retained after 30 min compared to the baseline.

### In vivo CFTR activity by Nasal Potential Difference (NPD)

+Rats were anesthetized with a cocktail of ketamine/dexmedetomidine (100mg/kg; 0.25mg/kg) by intraperitoneal injection. Under conscious sedation, nasal CFTR function was evaluated in rat and ferret nasal epithelium by previously described NPD protocol [20]. Briefly, a nasal catheter comprised of PE10 tubing was inserted into a single naris of an anesthetized rat or ferret for sequential infusion of Ringers solution (baseline); Ringers + Amiloride (100μM); Chloride-free Ringers solution consisting of KHPO (2.4mM), KHPO (0.4mM), Na Gluconate (115mM), NaHCO (25mM), and Ca Gluconate (1.24mM); Chloride-free Ringers + forskolin (20μM); and Chloride-free Ringers with 10μM of GlyH101. Chloride-mediated ion transport by CFTR was estimated as the mean changes in voltage in response to introduced solution infusions. Rats were allowed to recover from anesthesia by atipamezole (1mg/kg) administration for reversal and recovery.

### Tracheal CFTR assay by Ussing Chamber measurement

Chloride ion transport by CFTR channels in rat tracheal explants was estimated in short-circuit current (Isc) units with modified Ussing chambers (Physiologic Instruments, San Diego, CA) under voltage clamp conditions as previously reported [6, 7]. Rat tracheal segments were bathed in identical Ringer’s solution with 95% O_2_: 5% CO_2_ before sequentially adding apical Amiloride (100μM), apical low chloride Ringers, apical and basal forskolin (20μM), and apical CFTR-inh172 (10μM) (cells) or GlyH101 (10 μM) (trachea). Acquire and Analyze software was used to collect Ussing Chamber data monitoring epithelial ion transport Isc changes upon adding pharmacologic agents known to modulate CFTR activity.

### Mucociliary clearance apparatus by μOCT image analysis

Freshly excised rat and ferret trachea were imaged with a high-resolution reflectance imaging modality known as micro optical coherence tomography (μOCT), as previously described [8, 9]. Briefly, surface images from 8 different locations were acquired with an optical beam (Photonics Superk Extreme high-power supercontinuum White Light Laser, NKT Photonics) scanning longitudinally along the tracheal ventral surface with the larynx end serving as reference. Airway surface liquid (ASL) and periciliary liquid (PCL) depths were measured by ImageJ geometric tools. Mucociliary transport (MCT) rate was evaluated by tracking naturally occurring mucus particles traveling along the ventral surface over multiple frames, and then converting pixel to micron through the initial image calibrations [10].

### Statistics

Descriptive statistics were compared using Student’s t-test. Error bars represent mean ± SEM unless otherwise noted. All analysis parameters were set as two-sided and with α set to 0.05 to determine significance within GraphPad Prism (La Jolla, CA). SPSS ver. 25 (IBM, Armonk, NY)

## RESULTS

### Chronic alcohol administration in rats

The use of ethanol-containing liquid diets has been a well-established model in the alcohol field for the last 4 decades. Administration of alcohol in liquid diets causes no distress to the animals and no surgical procedures are required with this dietary feeding regimen.[11] Male and female 8-week-old rats expressing wild-type CFTR were pair-fed with alcohol or a control diet for 8 weeks. There are differences in the systemic levels of alcohol in different species due to pharmacokinetic differences. To control for such differences, we have developed a robust analytical chemistry approach to determine the alcohol burden in each rat enrolled in our studies by measuring ethanol metabolites. These data are also important to ensure that laboratory modeling of human alcohol consumption is physiologically valid and represents individuals with alcohol use disorders. Ethyl glucuronide (EtG) is a Phase II liver metabolite of ethanol used by forensic laboratories to estimate alcohol intake using blood/urine samples over a long time. Hence, to accurately model systemic levels of alcohol reached in our rat studies, we estimated EtG in blood collected 18 hr from the last feeding. As shown in **Supplementary Fig. 1** LC/MS/MS assay found a significantly increased EtG (Con: 111.1 ± 46, Alcohol: 2810 ±1120 ng/ml) in rat serum of alcohol when compared to their pair-fed controls. To minimize any species-specific metabolic differences that may restrict the reliable comparison of rat alcohol data with human databases, we quantified another common ethanol metabolite Ethylsulphate (EtS). Forensic analysts consider EtS to be a highly reliable biomarker for secondary validation. Our analysis of rat serum suggested increased EtS in alcohol-fed rats similar to that of EtG, suggesting any of these biomarkers can be used to reliably predict *in vivo* levels achieved in our animal studies. Based on the comparison between published data from multiple rodent and human studies, our chronic alcohol diet produced blood alcohol concentrations of ∼200 mg/dL. This alcohol dose agrees with preclinical alcohol studies from several in alcohol research.

### Delayed Mucociliary Clearance in alcohol-fed rats

Previous research indicates that alcohol administration significantly reduces ciliary beating in human cells and animal tissues. Alcohol exposure causes a time-dependent biphasic effect on ciliary beat frequency (CBF). Acute alcohol exposure up to 6 hours increases CBF [12, 13], whereas exposures beyond 6 hours significantly suppress CBF. [14, 15] These *in vitro* findings are supported by decreased CBF in rats administered with chronic alcohol [16]. Furthermore, alcohol was reported to cause an 8-fold increase in trachea-bronchial mucin (TBM) genes. [17] Thus, alcohol has the ability to modulate the formation of the airway mucus layer as well as ciliary activity.

Here, to determine if alcohol feeding in rats causes changes in whole lung MCC, we adapted a radiographic MCC assay routinely used in humans that relies on clearance of nebulized Technetium labeled radioparticles. Under sedation, Technetium particles were delivered directly into the lungs, and clearance was dynamically imaged every 2 minutes for the next 30 minutes. Rats fed with an alcoholic liquid diet exhibited a 42% (%clearance in 30 min, Con: 39.1 ± 4.8, Alcohol: 22.7 ±4.4) absolute reduction in MCC compared to iso-caloric controls (**Fig. 1**). Reductions in MCC were evident in all rats fed with alcohol diet, and this loss may not be due to acute effects of ethanol as the assay was conducted 18hrs from the last feed. There weren’t any differences in baseline deposition of radiotracer and MCC rates in both treatment groups were independent of gender.

**Figure 1.**
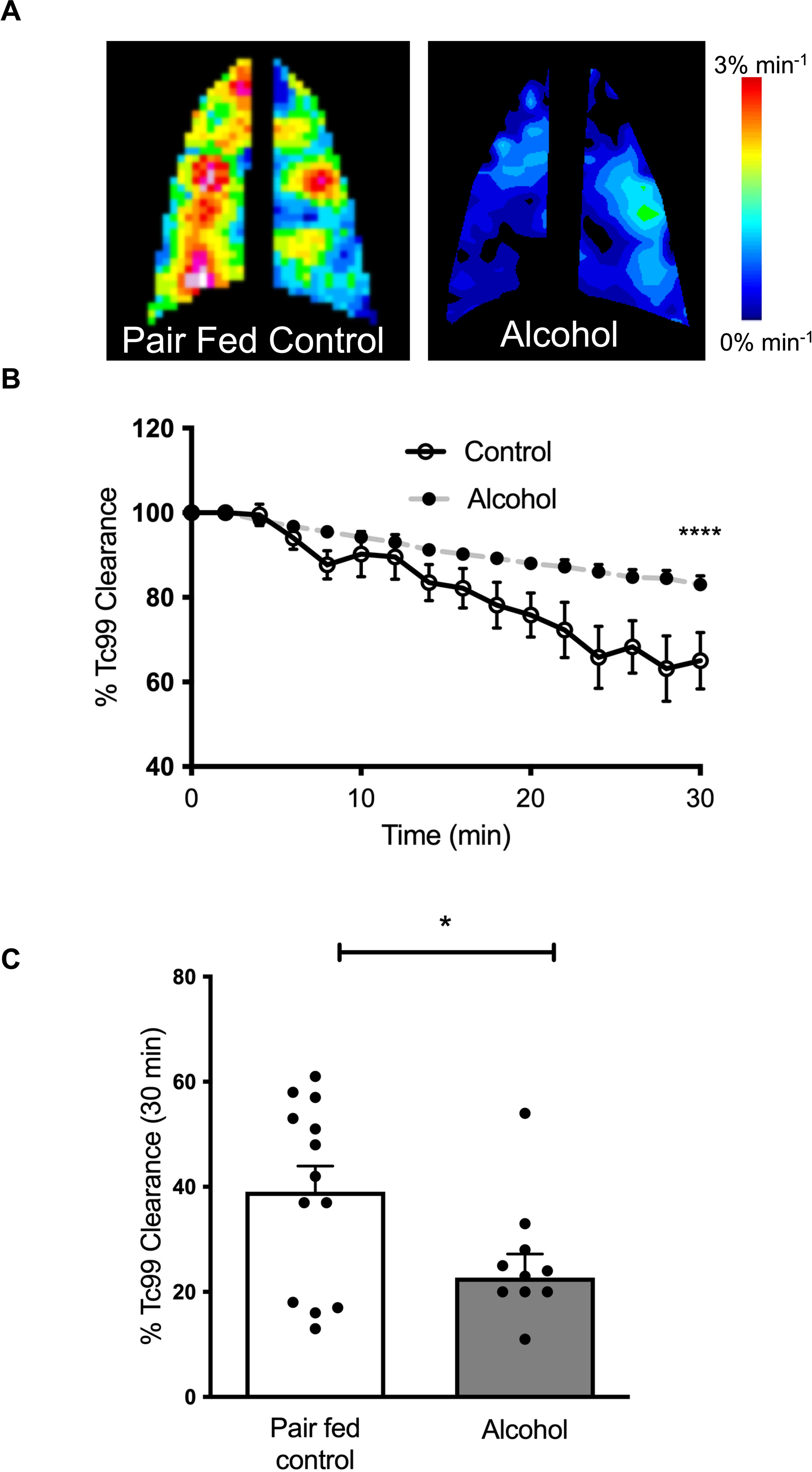
Alcohol reduces airway mucociliary clearance *in vivo*. Wild-type rats were administered either Lieber-DeCarli alcohol or an isocaloric control diet for 8 weeks. Mucociliary clearance (MCC) rate was estimated by tracking the elimination of inhaled Technetium Tc-99m DTPA (Tc99) from the lungs in rats under conscious sedation. (**A**) Reconstructed heat maps of rat lungs are shown, representing a range of clearance rates in 30 minutes. Red-maximal clearance of 3%/min, and blue/violet-minimal clearance approaching 0%/min. (**B**) Average percent clearance of Tc99 radiotracer over time is shown. (**C**) Summary graph shows the total percentage of radioparticles eliminated in 30 minutes in each animal. Statistical significance was assessed by an unpaired, 2-tailed non-parametric test (α=0.05), and error bars represent SEM. n=11-13/condition, *p<0.05, ****p<0.00005.

### CFTR Dysfunction in alcohol-administered rats

CFTR ion transport activity is critical for optical mucus hydration and clearance. Previously, we showed that alcohol exposure significantly reduced CFTR activity in bronchial epithelial cells treated with alcohol [3]. In a clinical report involving patients with alcohol-induced pancreatitis, that were similar decrements in CFTR activity. Hence, we measured *in vivo* CFTR activity in rats using nasal potential difference (NPD) assays that are standard in CF clinical diagnosis (**Fig. 2 A**). Alcohol administration caused a significant decrease in CFTR-mediated ion transport across the nasal epithelium of rats (**Fig. 2 C**). The magnitude of CFTR dysfunction among alcohol-treated rats was nearly 35% (CFTR function mV, Alcohol: –13.7 ± 1.0, Control: –21.0 ± 2.4 P≤0.01), matching what we previously reported in patients with mild inherited CF lung disease and also in those patients with smoking-induced COPD but without any genetic mutations in both CFTR alleles[6, 18]. Negative effects of alcohol on epithelial ion transport were very specific to CFTR channels as amiloride-sensitive Na+ absorption via ENaC channels was unaffected (**Fig. 5A, B**). Thus, these results indicate that clinically relevant CFTR dysfunction is present in the lungs of rats on an alcohol diet supporting further examination of its physiologic significance and impact on lung mucociliary defense. Just as importantly, the harmful effects of alcohol administration on CFTR function were also apparent when tracheal segments were analyzed *ex vivo* in modified-Ussing chambers. Trachea from alcohol-fed rats exhibited decrements in response to CFTR activator, forskolin, but not to amiloride, confirming the specific loss of CFTR but not ENaC function in rat airways **(Fig 3 A, B**). These differences in epithelial ion channel activity were not associated with decrements in transepithelial electric resistance, suggesting CFTR loss is unrelated to epithelial barrier leak (**Fig 3 C**). Next, we examined the alcohol effect on lung CFTR expression by quantifying absolute mRNA abundance by droplet digital PCR (**Fig 3 D**) and Western Blot protein analysis (**Fig 3 E**). Interestingly, there was no apparent difference in CFTR mRNA abundance. However, the representative blot illustrates diminished immature B band compared to β-actin, whereas the membrane surface-bound CFTR C band remained unaffected (**Fig 3 F, G**). Thus, it is likely that changes in lung CFTR expression may not fully explain alcohol induced CFTR dysfunction indicating the need for more exhaustive research into the underlying mechanisms.

**Figure 2.**
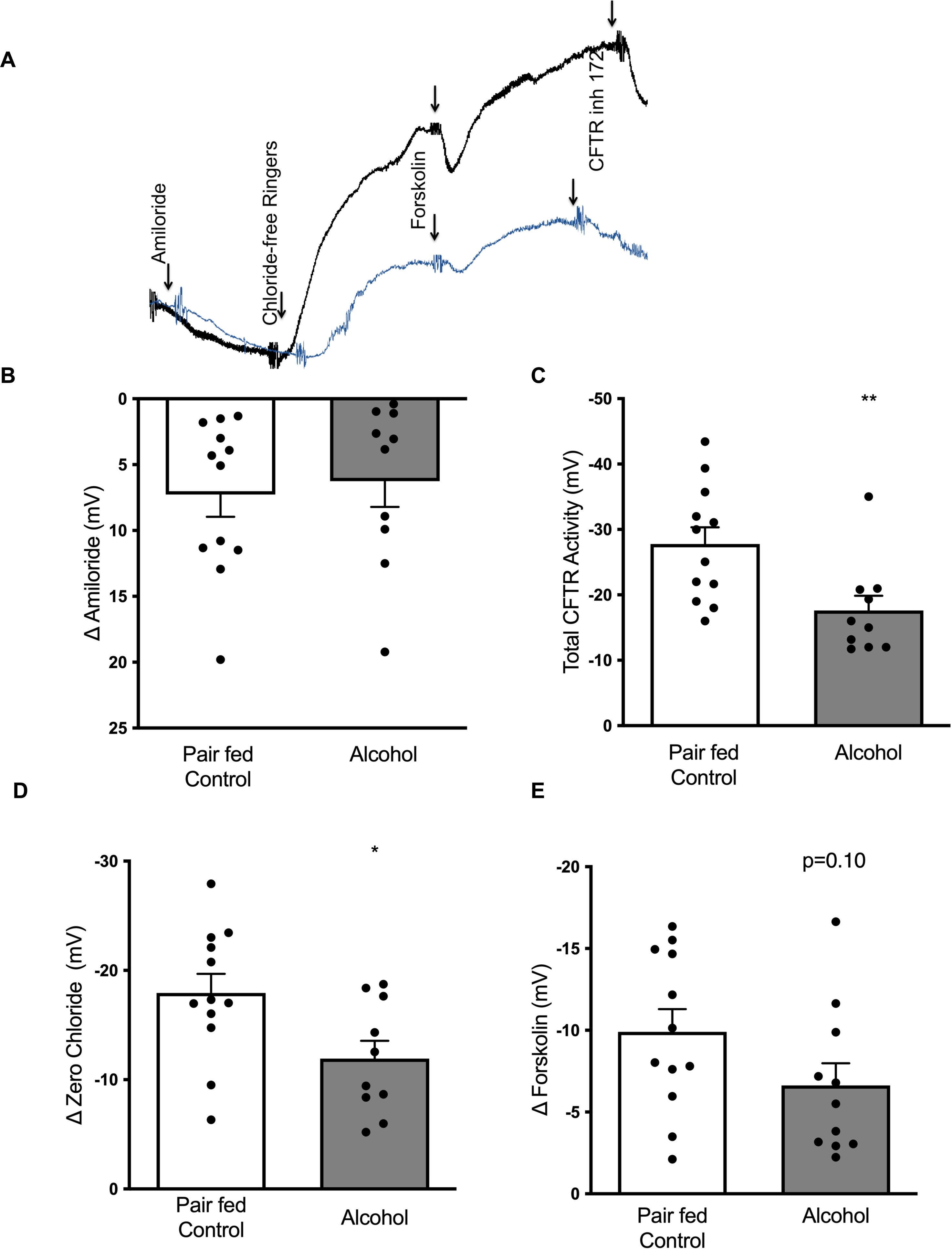
Alcohol reduces airway CFTR function but not ENaC. *In vivo,* Epithelial ion transport was assessed by nasal potential difference (NPD) assay in rats expressing CFTR(+/+) under conscious sedation. (**A**) Representative tracings of voltage change (mV) across nasal epithelium in wild-type rats fed with Lieber-DeCarli alcohol (**blue**) or isocaloric control (**black**) diet for 8 weeks are shown. Summary bar graphs illustrate that alcohol intake does not alter ENaC activity (**B**) but significantly reduces total CFTR activity (**C**). When nasal CFTR activity was differentiated based on gating status at baseline, the adverse impact of alcohol was observed in the portion of channels that were open (**D**) and closed (**E**). Data were averaged and expressed as mean ± SEM. Statistical significance was assessed by an unpaired, 2-tailed non-parametric test (α=0.05). n=10-12/group, *p<0.05, **p<0.005.

**Figure 3.**
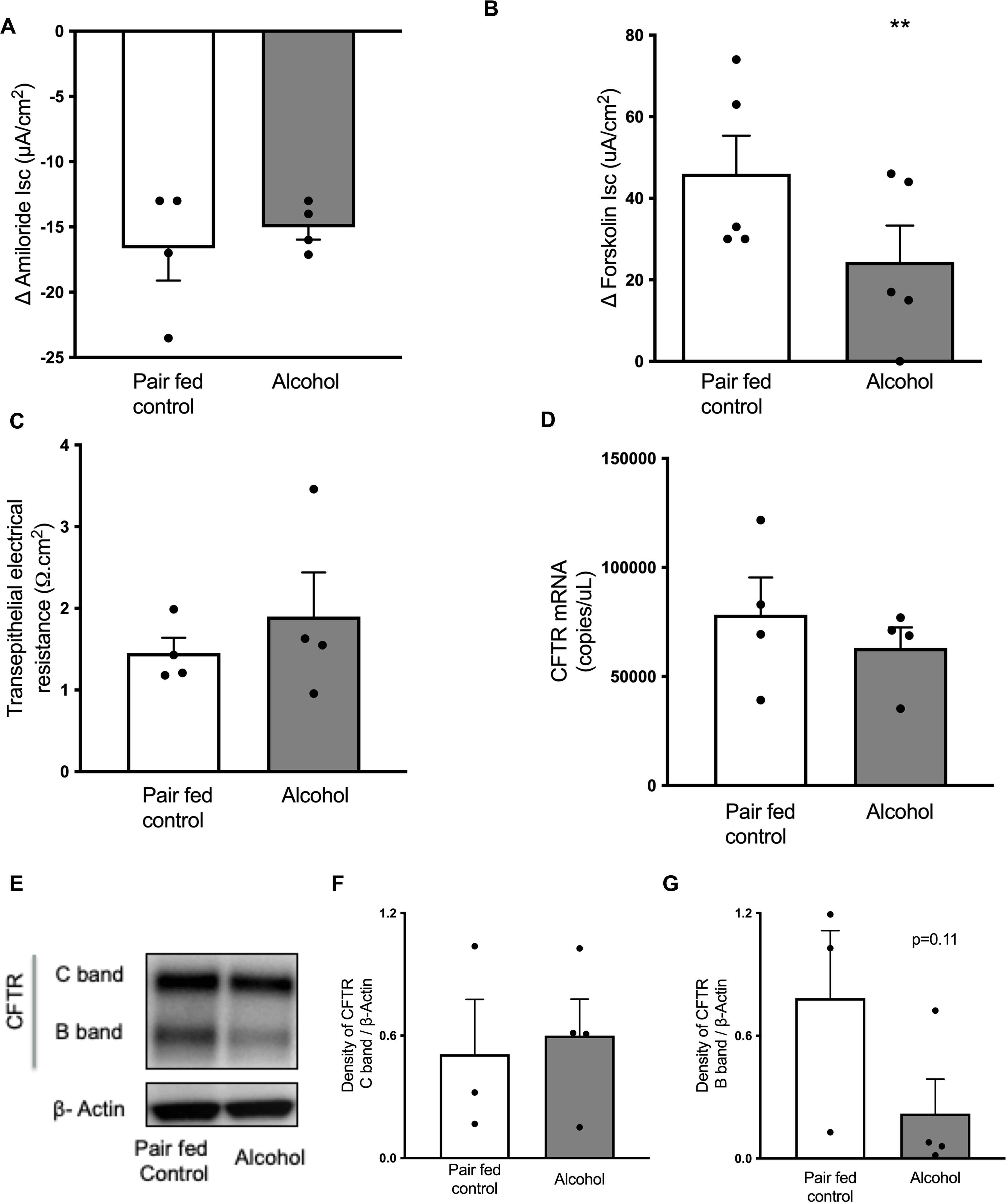
Alcohol reduces conducting airway CFTR function and expression. Freshly excised trachea from rats expressing CFTR (+/+) and fed 8 weeks with lieber-decarli alcohol or isocaloric control diet were mounted into Ussing chambers to assess short circuit current (Isc) changes in airway epithelial ion transport. Summary graphs illustrate Isc changes representative of amiloride-sensitive sodium absorption (**A**) and forskolin-stimulated anion transport via CFTR channels (**B**). These differences in epithelial ion channel activity were not associated with decrements in transepithelial electric resistance (Ω.cm^2^) (**C**). Lung CFTR expression was quantified by absolute mRNA abundance by droplet digital PCR (**D**) and Western Blot protein analysis (**E**). The representative blot illustrates the membrane surface-bound CFTR C band and the immature B band as compared to β-actin. (**F, G**) Summary graphs quantify the ratio between the density of the C band/β-actin and the density of the B band /β-actin. Data were averaged and expressed as mean ± SEM with statistical significance assessed by unpaired, 2-tailed non-parametric test (α=0.05) using GraphPad PRISM (GraphPad Software Inc.).(A-D n=5/condition; F-G n= 3-4/condition; G n=4/condition; *p<0.05, **= p<0.005)

### Alcohol impairs airway surface hydration and mucus transport

To verify if alcohol effects on CFTR resulted in downstream alterations in airway epithelial functions relevant to MCC, we conducted µOCT imaging of freshly harvested rat trachea (**Fig. 4A)**. These images were used to quantify changes in airway surface hydration by measuring the height of the PCL layer (liquid sheath within ASL most critical for MCC). As shown in **Fig. 4**, alcohol administration significantly reduced PCL height (**Fig. 4C)**, and mucus transport (MCT) rate (**Fig. 4D)**, consistent with *in vivo* MCC data (**Fig. 1**). However, in alcohol-treated rats ASL (**Fig. 4B)** appeared to tend towards a slight increase, suggesting an increased production of mucus. Histopathological analysis of rat lungs stained with AB-PAS staining revealed mucus hypersecretion and goblet cell hyperplasia with alcohol administration (**Fig. 5A, B**). Since Muc5b is the most important mucin for maintaining MCC and effective bacterial clearance (Muc5ac was found to be dispensable in rodents),[19] we measured its distribution using a specific antibody in rat lungs. Alcohol administration significantly elevated the production of Muc5b and more importantly (**Fig. 5C, D**). Collectively, we find reduced CFTR-mediated ion transport by alcohol results in a mucus expression: fluid secretion imbalance severely inhibiting mucus transport.

**Figure 4.**
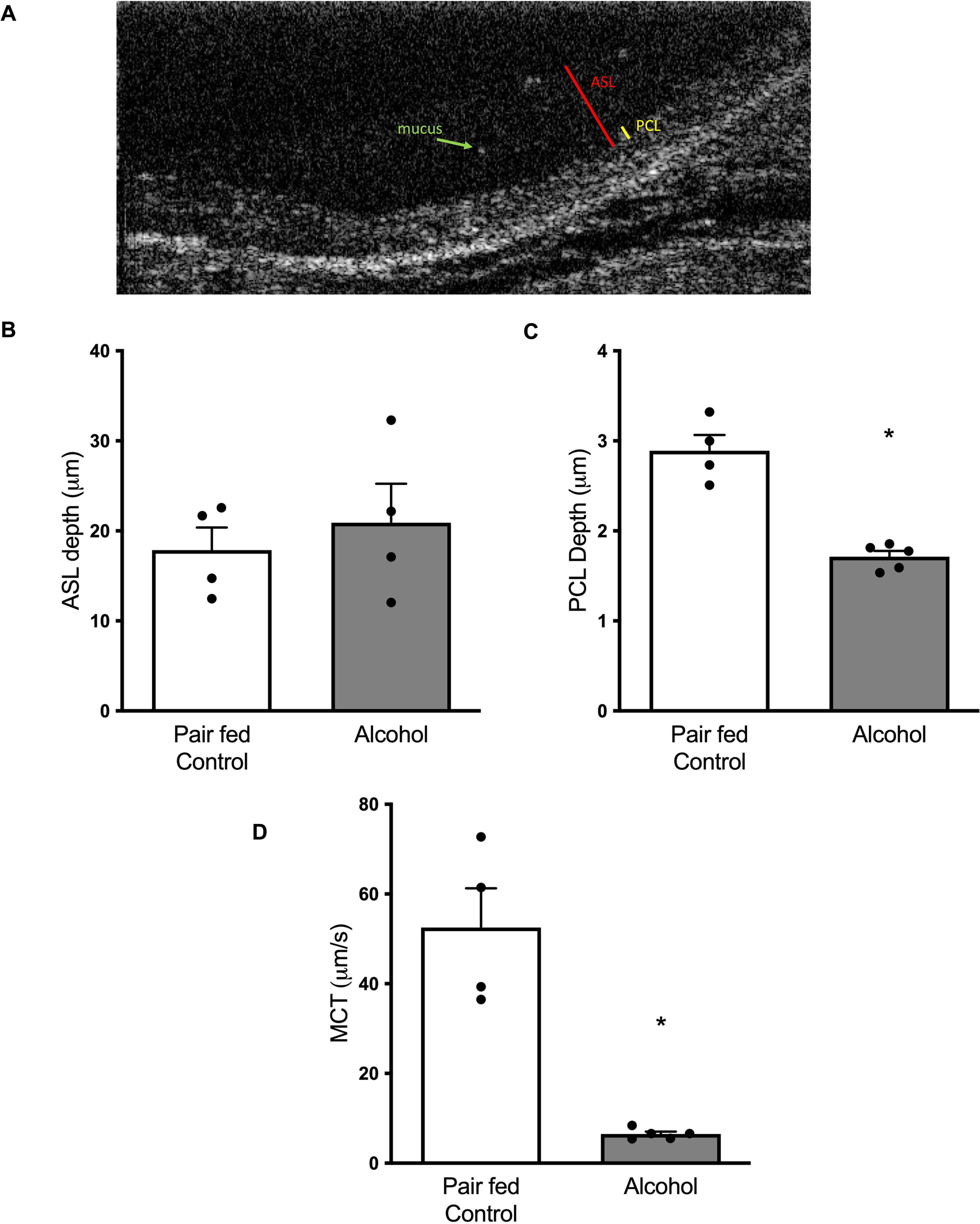
Alcohol consumption impairs airway surface hydration and mucus transport in rats. Freshly excised trachea from rats that were fed either lieber-decarli alcohol or isocaloric control diet for 8 weeks were imaged by micro optical coherence. tomography (μOCT) to assess physiologic changes in the airway mucociliary clearance parameters. (**A**) Representative tracheal mucosal surface image indicating air surface liquid (ASL) depth (μM), denoted in red and summarized in **B**, and periciliary liquid layer (PCL) depth (μM), shown in yellow and summarized in **C**. (**D**) Mucociliary transport (MCT) rates (μM/s) were calculated by tracking native mucus particles, an example indicated by a green arrow, across multiple regions of interest along the ventral surface of the trachea. Data were averaged and expressed as mean ± SEM with statistical significance assessed by unpaired, 2-tailed non-parametric test (α =0.05) using GraphPad PRISM (GraphPad Software Inc.). (n=4-5 rats/condition; *=p<0.05; **= p<0.005)

**Figure 5.**
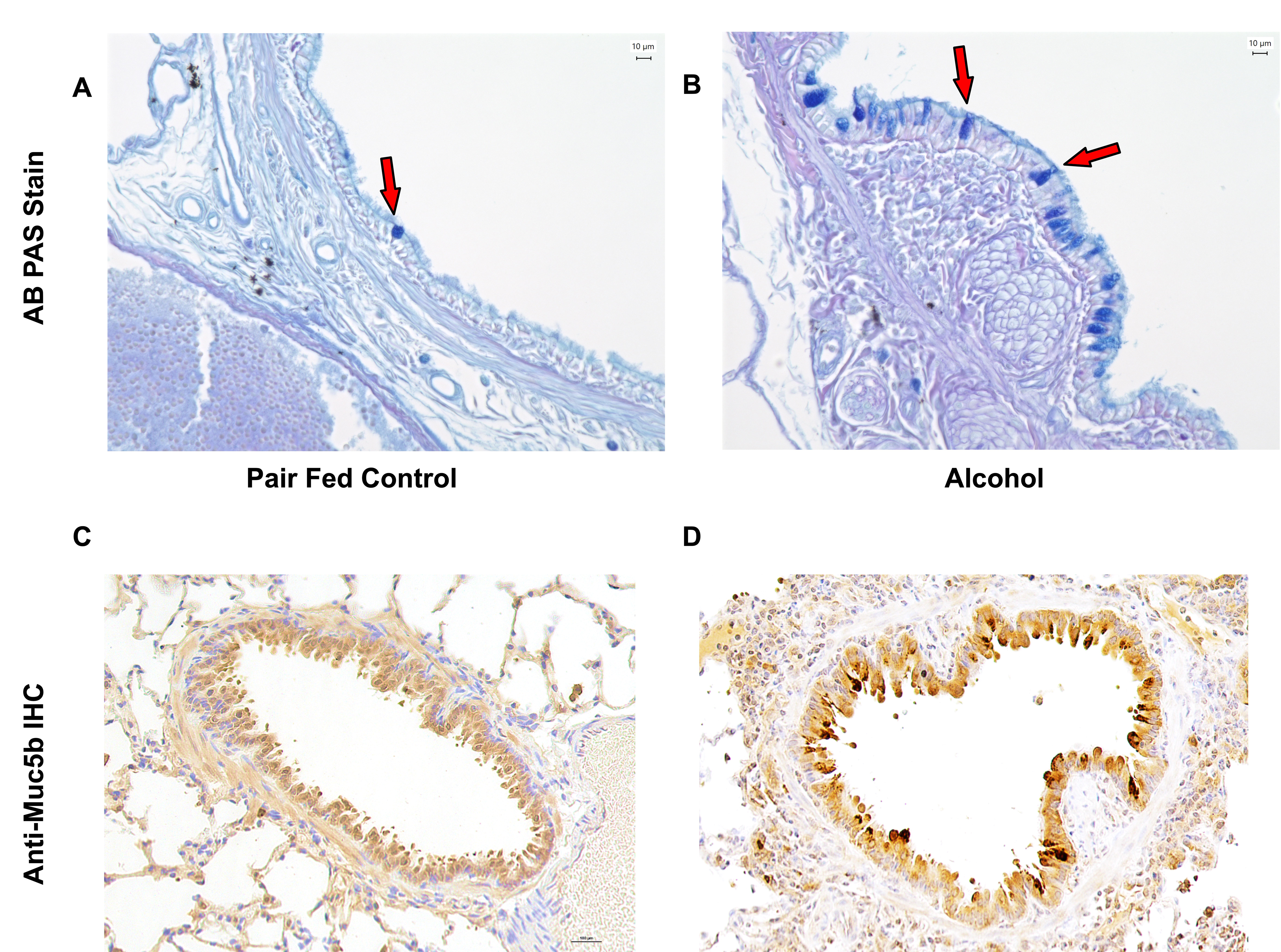
Alcohol diet promotes goblet cell hyperplasia and mucus overproduction in rat airways. Lung sections from wild-type rats fed with 8 weeks of lieber-decarli alcohol or isocaloric control diet were analyzed for mucus expression. (**A, B**) Representative 20X images of AB-PAS staining of rat airways indicate alcoholic diet increases the number of mucus-abundant goblet cells (red arrows) and (**C, D**) immunohistochemical staining with a specific antibody against MUC5b.

### CFTR dysfunction by alcohol is acquired and unaltered by baseline CFTR

To determine the pathophysiologic and clinical impacts of heavy drinking in carriers of genetic CFTR mutations on one allele (mutations on both alleles would qualify CF diagnosis), we conducted alcohol studies in rats with heterologous CFTR (Het, CFTR +/-). These studies enabled us to determine gene-dose responses on alcohol effects and to distinguish the contribution of genetics and environment to acquired dysfunction. Data obtained with these heterozygote rats was compared to that of wild-type rats treated with alcohol and wild-type rats treated with a control diet. Het rats represent CF carriers (parents of CF patients) known to suffer higher rates of respiratory infections and are known to express just one functional CFTR allele, nearly 50% of lower CFTR protein expression. Not surprisingly, even with a control diet, MCC rate and CFTR function were lower in Het rats compared to wild-type counterparts (**Fig. 6 A, B**). Interestingly, proportions of alcohol-induced defects in CFTR and MCC rates were similar in both genotypes suggesting alcohol indiscriminately disrupts airway epithelial functions. Thus, these results provide compelling genetic evidence for the role of acquired CFTR dysfunction in alcohol-induced decrements in whole lung MCC. Similarly, CFTR genotype had no bearing on alcohol effects on ENaC function, validating the specificity of the pathogenic mechanism in rats.

**Figure 6.**
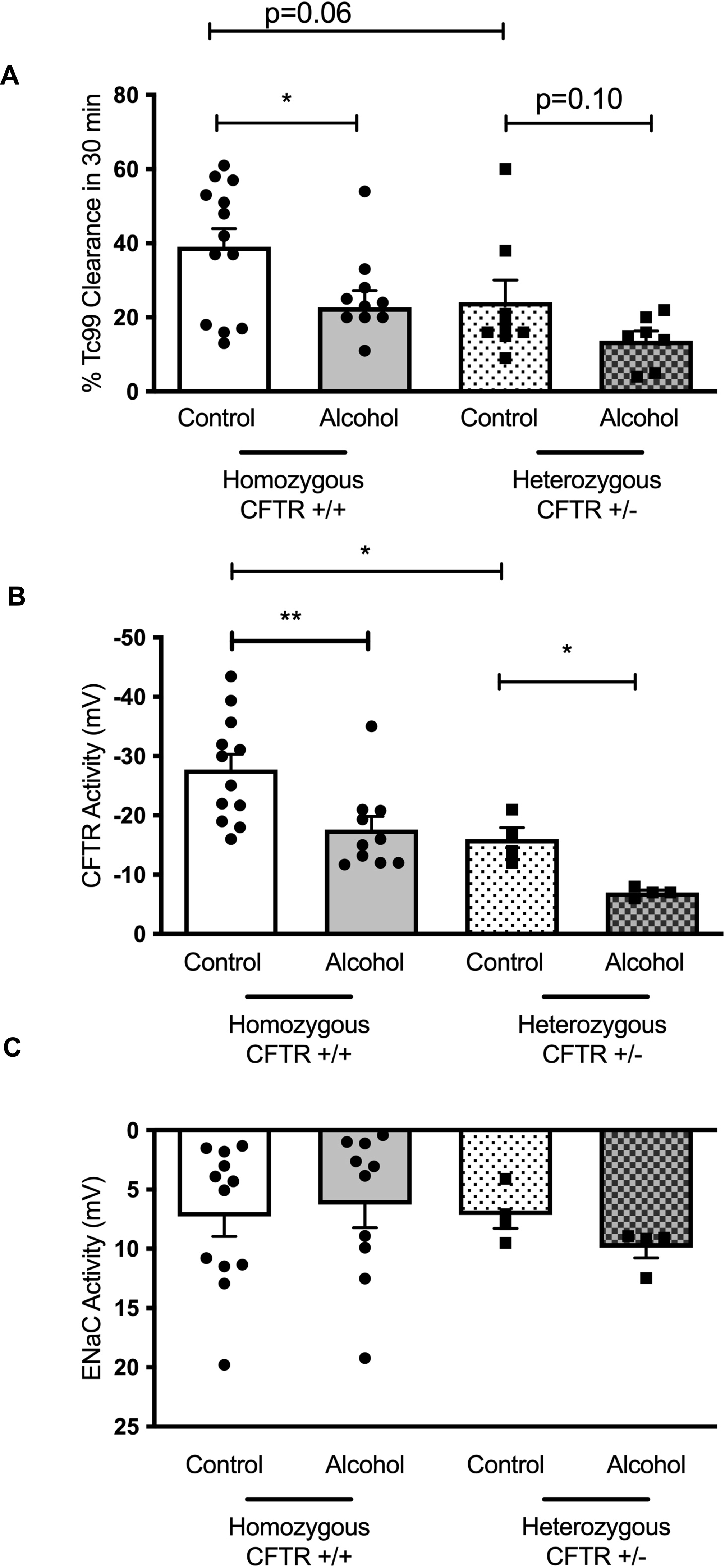
Alcohol impairment of CFTR function and Mucus Clearance in rats is independent of CFTR genotype. Data illustrate alcohol indiscriminately disrupts airway epithelial function in rats homozygous for CFTR (+/+) or heterozygous for CFTR (+/-) following 8 weeks of administration with a lieber-decarli alcohol diet. (**A**) Alcohol delayed the clearance of DTPA-conjugated technetium-99 (Tc99) from the lungs of rats expressing CFTR+/+ and CFTR+/-. (**B, C**) ENaC and CFTR function were measured in rats homozygous and heterozygous for CFTR by in vivo nasal potential difference (NPD) assay. Data from homozygous (n=10-12/condition) and heterozygous (n=4/condition) CFTR-expressing rats were averaged and expressed as mean ± SEM. Statistical significance was assessed by an unpaired, 2-tailed non-parametric test (α=0.05) using GraphPad PRISM (GraphPad Software Inc.). (*=p<0.05; **=p<0.05).

## DISCUSSION

Mucociliary clearance is the primary defense mechanism of airways against inhaled pathogens, irritants and allergens. Association of alcohol use with impaired MCC is known for decades.[20] When MCC is delayed, the retention time of pathogens and irritants is increased leading to enhanced pathogenicity.[21] Mucociliary dysfunction is also a common feature of chronic airway diseases such as cystic fibrosis, primary ciliary dyskinesia, asthma, and chronic bronchitis with recurrent lung infections.[22] More importantly, drugs and other interventions that successfully augment MCC offer significant clinical benefits by relieving airway obstruction and by reducing infectious exacerbations.[23, 24] Thus, restoration of diminished MCC to full physiologic levels represents a great potential to rapidly remove inhaled/aspirated pathogens are prevent pathogenic lung colonization in excessive alcohol use prone to frequent infections. Hence, understanding the role of CFTR dysfunction in alcohol-use disorders is highly relevant.

Data presented in this report demonstrated for the first time the magnitude and dose dependence of alcohol-induced CFTR dysfunction in an animal model of chronic alcohol administration.[2] These data successfully verified alcohol effects in primary human bronchial epithelial (HBE) cells derived from non-CF (CFTR +/+) donors and also tested if the observed CFTR defects were only in response to adenosine or were broadly applicable to all physiologic regulators by using forskolin, the most common CFTR agonist. While cigarette smoking, another common agent that is known to affect both CFTR and ENaC channels, alcohol effects are CFTR-specific.[25, 26] It’s also worth noting that CFTR activity measures across nasal epithelium were reflective of ion transport status in lower airway epithelium in rats and were in agreement with prior studies in mice[7], rats[27], ferrets[28] and human smokers and COPD patients.[25] This congruency of alcohol-induced CFTR defects is also reflected in CFTR activity measures in tracheal explants from the most recent cohort of alcohol-fed rats (**Fig. 3**). Thus, these *in vivo* results validate clinically relevant CFTR dysfunction supporting a further examination of physiologic significance and impact on lung antimicrobial defense.

While there haven’t been any clinical reports on CFTR function in patients with alcohol-related lung disease, Maleth *et al* published reduced CFTR activity in patients with alcohol-induced pancreatitis. In healthy volunteers (N= 49) with acute alcohol consumption exhibited increased sweat chloride levels (suggesting reduced CFTR function) that return to normal levels upon sobriety. However, chronic alcohol consumption had a lasting negative effect on CFTR activity, as found in patients with a history of chronic alcohol use (N= 15) despite abstinence.[3] Interestingly, expression of total CFTR mRNA and protein in pancreatic duct epithelial cells were not reduced in these patients with alcohol-induced chronic pancreatitis. This is consistent with our own findings of unaltered CFTR mRNA and partial change in CFTR band B that is not know to have ion transport function protein expression (**Fig 3)** in the lungs of alcohol-administered rats despite, significant decreases in ion transport activity. This contradiction prompted an editorial that concluded that underlying causes for alcohol-reduced CFTR function may not be directly related to protein levels but, to altered protein folding and regulation on the cell surface.[29].

Previously, we evaluated the impact of alcohol exposure on the ability of epithelial cells to generate cAMP levels required for CFTR phosphorylation and channel gating. Compared to control cells, adenosine addition (as opposed to forskolin used in this study) for 10 minutes, the time point at which the epithelial CFTR currents were at their peak, elicited lower intracellular cAMP generation in Calu-3 cells treated with alcohol. To distinguish if the decrease in cAMP was in fact the basis for CFTR dysfunction by alcohol, Sp-cAMPS, a cell permeable synthetic analogue of cAMP, was supplemented to alcohol-treated cells. 10 µM Sp-cAMPS completely eliminated suppressive effects of alcohol on adenosine-induced CFTR activity (**Fig. 11B**). These data are consistent with prior alcohol studies that documented alcohol exposure decreases cAMP signaling and protein kinase A (PKA) enzymatic activity [12, 30]. Interestingly, alcohol reduction of cAMP was not due to its inhibition of adenylyl cyclases that generate cAMP but likely due to increased activity enzymes that specifically degrade cAMP.

Mechanistically, we were able to link CFTR defects to PCL dehydration, explaining altered mucus clearance rate in vivo and in vitro. That said, reduced PCL depth may not fully explain the entirety of the dramatic decline in mucus transport rate, implying additional CFTR-dependent abnormalities in mucus properties in alcohol-fed rats, such as viscoelastic properties likely from more abundant mucus secretion by alcohol. Thus, additional research is needed to verify if alcohol exposure increases the viscoelasticity of airway mucus, contributing to delayed clearance. This notion is informed by mucus hypersecretion and goblet cell hyperplasia with alcohol administration (**Fig. 5**). Since Muc5b is the most important mucin for maintaining MCC and effective bacterial clearance (Muc5ac was found to be dispensable in rodents),[19] we measured its distribution using a specific antibody in rat lungs. Alcohol administration significantly elevated the production of Muc5b compared to their age-matched healthy controls.

Collectively, we find reduced CFTR-mediated ion transport by alcohol results in a mucus expression: fluid secretion imbalance severely inhibiting mucus transport. Since alcohol is known for its pleiotropic effects, we compared the alcohol effects on CFTR activity and MCC rate to CFTR heterozygote (Het, CFTR +/-) rats treated with an isocaloric control diet for the same duration. Het rats express just one functional CFTR allele and represent CF carriers (parents of CF patients) known to suffer higher rates of respiratory infections. Alcohol-induced defects in CFTR and MCC rate approximate in vivo findings in Het rats. Thus, these results provide compelling genetic evidence for the role of acquired CFTR dysfunction in alcohol-induced decrements in lung mucociliary clearance defense and compromised lung defense in alcohol abusers. Data obtained from these studies support that CFTR may be a suitable target for treating recurrent lung infections in individuals with a history of alcohol disuse using CFTR modulator drugs that are in clinical use for patients with cystic fibrosis and various stages of clinical testing for acquired CFTR defects in smokers with COPD and infectious exacerbations.

