## Supplementary Fig. 1 for "Alcohol-Induced Mucociliary Dysfunction: Role of Defective CFTR Channel Function"

### Supplementary Figure 1

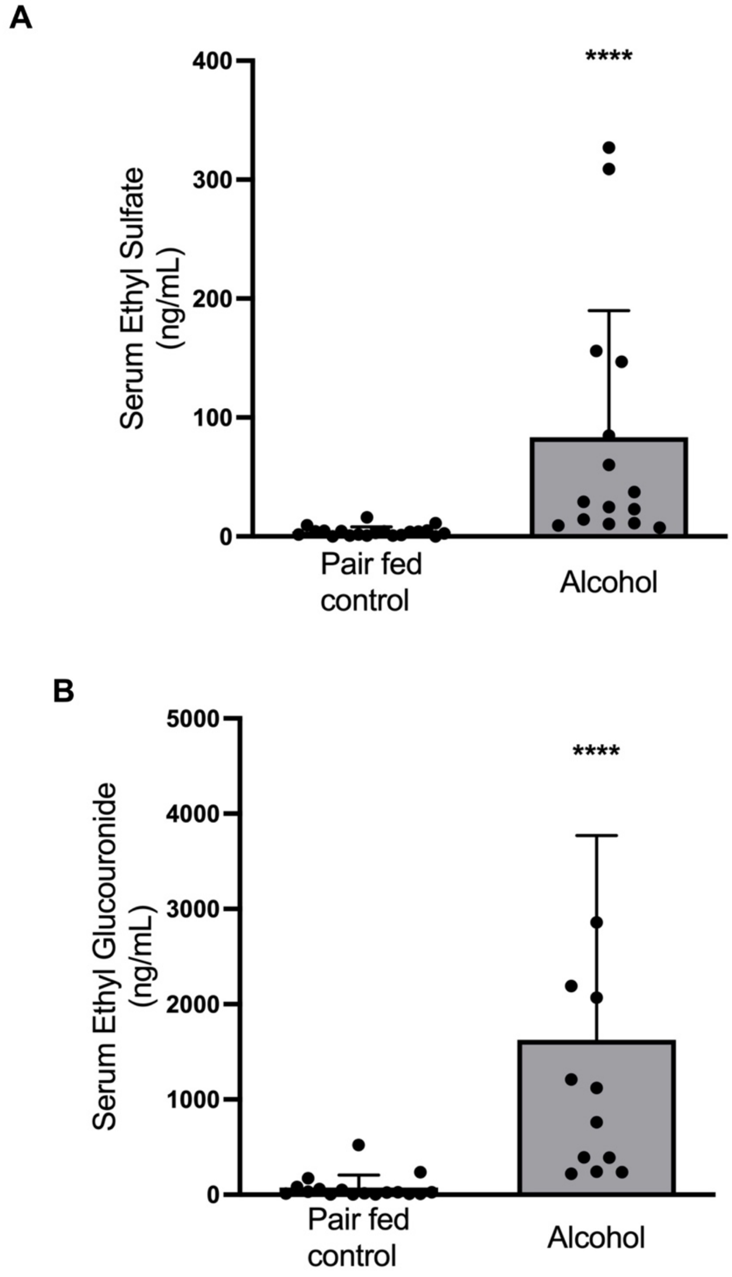


**Supplement 1. Blood alcohol levels in alcohol-fed rats.** Summary bar graphs demonstrate stable ethanol metabolites of ethyl sulfate (**A**) and ethyl glucuronide (**B**) in the serum from Sprague-Dawley rats expressing wild-type CFTR (+/+) fed for 8 weeks with either lieber-decarli alcohol or isocaloric control diet. Values were averaged and expressed as mean ± SEM. Statistical significance was assessed by an unpaired, 2-tailed non-parametric test ($\alpha$=0.05) using GraphPad PRISM (GraphPad Software Inc.). (A-B n=10-14/condition; ****p<0.0005).
